# A Hidden Markov Model for Investigating Recent Positive Selection through Haplotype Structure

**DOI:** 10.1101/011247

**Authors:** Hua Chen, Jody Hey, Montgomery Slatkin

## Abstract

Recent positive selection can increase the frequency of an advantageous mutant rapidly enough that a relatively long ancestral haplotype will be remained intact around it. We present a hidden Markov model (HMM) to identify such haplotype structures. With HMM identified haplotype structures, a population genetic model for the extent of ancestral haplotypes is then adopted for parameter inference of the selection intensity and the allele age. Simulations show that this method can detect selection under a wide range of conditions and has higher power than the existing frequency spectrum-based method. In addition, it provides good estimate of the selection coefficients and allele ages for strong selection. The method analyzes large data sets in a reasonable amount of running time. This method is applied to HapMap III data for a genome scan, and identifies a list of candidate regions putatively under recent positive selection. It is also applied to several genes known to be under recent positive selection, including the *LCT, KITLG* and *TYRP1* genes in Northern Europeans, and *OCA2* in East Asians, to estimate their allele ages and selection coefficients.

## 1. Introduction

Natural selection plays an important role in the recent history of human evolution, and is still active in shaping the genetic diversity pattern of human populations. Genes under positive selection may be involved in the adaption to new environments and in the resistance to infectious diseases (Hamblin et al., 2002; Bersaglieri et al., 2004; Tishkoff et al., 2007; Simonson et al., 2010; Yi et al., 2010; Beall et al., 2010; Peng et al., 2011; Xu et al., 2011; Xiang et al., 2013). In recent years, interest is growing in detecting positive selection using DNA polymorphism data, since the rapid accumulation of genomic level molecular polymorphism data provides a chance to systemically investigate the footprints of natural selection (Tajima, 1989; Fu and Li, 1993; Fay and Wu, 2000; Akey et al., 2002; Sabeti et al., 2002; Kim and Stephan, 2002; Nielsen et al., 2005; Voight et al., 2006; Tang et al., 2007; Sabeti et al., 2007; Williamson et al., 2007; Pickrell et al., 2009; Chen et al., 2010; Grossman et al., 2013). Recent positive selection (RPS), which occurred in the recent past and is still active, has gained particular attention. RPS can increase the frequency of advantageous alleles in a short time, and thus result in high level of haplotype sharing in the vicinity of the selected mutant, and higher homozygosity among the selected haplotypes than those carrying the neutral allele. This unique pattern of multilocus haplotype structure enables methodology development for identifying genes under RPS and parameter inference of the selection process.

Statistical tests have been developed to test for natural selection based on multilocus haplotype frequency distribution or haplotype structure (e.g., Ewens 1972; Slatkin 1994; Depaulis et al. 1998; Innan et al. 2005). Innan et al. (2005) presented a good review of these haplotype-based methods. Among the various haplotype-based tests, several exploit the specific haplotype structure caused by RPS by comparing the homozygosity level between selected and neutral haplotype groups (e.g., Hudson et al. 1994; Sabeti et al. 2002; Hanchard et al. 2005; Voight et al. 2006). The first of this kind was proposed by Hudson et al. (1994). Their haplotype test was designed to examine a group of high frequency haplotypes with little genetic variation among them. The test was carried out by estimating the probability of observing fewer polymorphic sites in repeated coalescent simulations given the sample size and allele counts. Hudson et al (1994) applied the method to analyze the *Sod* gene in *Drosophila melanogaster.* At this locus, there are two alleles, labelled by “slow” and “fast”. Hudson et al (1994) found that there was no mutation among the slow allele group, which has a frequency of approximately 18%, and concluded that there was a significant deviation from neutrality. Sabeti et al (2002) proposed a Relative Extended Haplotype Homozygosity (REHH) test, which starts with choosing a “core region” (Sabeti et al., 2002), a small region of very low historical recombination, and then calculates as the test statistic the ratio of Extended Haplotype Homozygosity (EHH) of the core haplotype under test over the other core haplotypes. The significance level of the REHH test is generated by coalescent simulation of neutral data that match to the real data by haplotype group numbers and polymorphism level. The method was applied to identify the selected core haplotypes in two malaria-resistance genes *G6PD* and *CD40*, and later to the HapMap data for a genome-wide scan (Sabeti et al., 2007). The iHS (integrated haplotype score) test, as a variant of REHH test, was proposed by Voight et al (2006). The integrated EHH (iHH, defined as the area under the EHH curve) for the ancestral and derived alleles of the mutant are first estimated. The iHS score is then standardized to follow a normal distribution approximately, and subsequently used to test the deviation from neutral model.

In addition to detecting selection, one may be also interested in estimating the selection intensity and the timing of the selection process. There are several methods for this purpose (Slatkin and Rannala, 1997; Slatkin, 2000; Slatkin and Rannala, 2000; Slatkin, 2001, 2002; Kim and Stephan, 2002; Coop and Griffiths, 2004; Rannala and Reeve, 2004; Slatkin, 2008; Chen and Slatkin, 2013). Among them, some consider a single marker linked to the selected locus (Slatkin and Rannala, 1997; Slatkin, 2000, 2001; Kim and Stephan, 2002); and only a few of them model the haplotype structure of multiple marker loci (Coop and Griffiths, 2004; Rannala and Reeve, 2004; Slatkin, 2008; Chen and Slatkin, 2013). Coop and Griffiths (2004) developed a full likelihood method under the structured-coalescent framework (Hudson and Kaplan, 1988). They adopted the time-reversible Moran model to first simulate the allele frequency trajectory of the selected mutant, and then conditioning on the trajectory, they were able to simulate the genealogical history of the sample. The limitation of their method is that only mutations among different haplotypes are considered and the method is only applicable to non-recombining regions. Rannala and Reeve (2003) modelled both recombination and mutation, but their method depicts the haplotype structure in the vicinity of mutants under neutrality and has unrealistic assumptions of constant allele frequencies for all loci during the selective process. Slatkin (2008) used a linear birth-and-death process to simulate the allelic genealogies of selected mutants and modelled the multilocus haplotype structure under the influence of both recombination and mutation. Chen and Slatkin (2013) also proposed a multilocus haplotype model that describes the dynamics of the haplotype structure under the joint effects of selection, recombination and mutation, by efficiently reducing the complexity of state spaces. Their method exploits the importance sampling approach to generate the historical allele frequency trajectory of the selected mutant, and thus works for populations with temporally changing size (Slatkin, 2001). All the methods are coalescent-based and take into account of randomness of trajectory and genealogies by Monte Carlo averaging, which requires intensive computation. In comparison to the above computationally intensive methods, Voight et al (2006)’s approach is simplified and computationally feasible. Their method estimates the distance at which the haplotype sharing decreases to a pre-chosen level, and then assumes the decaying of haplotype sharing follows a Poisson process. Voight et al (2006)’s method further assumes the independent histories of different haplotypes to avoid intensive computation due to the integration over unknown gene genealogies, and thus is suitable for whole-genome analysis.

In this paper, we propose a hidden Markov model to identify the ancestral haplotypes retained during the selective process for the purpose of both detecting selection and estimating the selection intensity. Comparing to the existing methods, e.g., the REHH and iHS tests, which use summary statistics to evaluate the similarity of haplotypes, our method is model-based so that it has the potential to be extended to more complicated scenarios, such as, multiple ancestral haplotype groups (soft sweeps on standing variation, Hermisson and Pennings 2005), haplotype data from multiple populations, and genotype data with unknown phase etc.

The method is also different from the aforementioned coalescent-based models in that we do not try to simulate gene genealogies among individuals and the events occurring along the genealogies by Markov Chain Monte Carlo or importance sampling approaches (Slatkin, 2008; Coop and Griffiths, 2004; Chen and Slatkin, 2013). Our method is similar to that of Voight et al. (2006) in this respect. We treat each haplotype independently by assuming a “star” genealogy and ignore the randomness of frequency trajectory of the selected allele. Both methods are computationally efficient and applicable to genome-wide analysis. Compared to Voight et al (2006), our method provides a better estimation of the selection coefficient when the selected allele is common or nearly fixed, since we explicitly model the probability of effective recombination causing the break of ancestral haplotype extents, which is different from the simple recombination process in Voight et al (2006) and others. As we will show in a later section, when the selected mutant is at high frequency, the bias in the Voight et al (2006) method can be as high as ≈ 20%.

The aim of this paper is twofold: first, we propose a hidden Markov model (HMM) that can explore the haplotype structure of a genomic region, and the inferred haplotype structure can be used to detect the existence of selection; second, we use a simplified population genetic model for the ancestral haplotype extent inferred from the HMM to estimate the selection intensity and the allele age. In the following sections, we first elucidate the details of the method. We then use coalescent simulations to investigate the power of detecting RPS and the accuracy of parameter estimation. We apply the method to analyze several well-known genes under RPS to demonstrate its performance, including the lactose persistence gene (*LCT*) in Northern Europeans, and *KITLG, TYRP1* and *OCA2*, known to confer skin pigmentation in Northern Europeans or East Asians.

## 2. Methods

In this section, we first present the HMM for identifying the extent of ancestral haplotypes. Two tests are further developed based on the HMM for detecting RPS. We then describe a population genetic model of hitchhiking. To be specific, we determine the allele frequency of a selected mutant and the approximate distribution of ancestral haplotype extents as a function of selection intensity and time, and then use this model to infer the selection intensity and the allele age of the selected mutant.

### 2.1. Data and parameters

The input data is a sample of *n* chromosomes (haplotypes) randomly collected from a contemporary population and genotyped at *m* SNP loci, and the phase of the chromosomes is assumed to be known. The data *X* is an *n* by *m* matrix, with the entry *X*_*i*,*j*_ encoded as 0 or 1 to denote the allele type of the *j*th SNP on the *i*th chromosome. The physical and genetic positions of the SNPs are also assumed to be known. The parameter set, Γ = {*s*,*t*, **A**, λ}, include the selection coefficient (*s*), the position of the advantageous mutant (λ), the mutant age (*t*), and the ancestral haplotype: **A** = (*A*_1_, *A*_2_,…, *A*_*m*_), which is not observable. Here *A*_*j*_ denotes the allele at the *j*th position of the ancestral haplotype. We use the term “ancestral haplotype” to refer to the alleles in every SNP position along the chromosome from which the advantageous mutant first arose. Since we assume a hard sweep model for RPS, there exists only one ancestral haplotype. But the assumption can be easily relaxed to model soft sweeps by allowing multiple ancestral haplotypes.

### 2.2. A hidden Markov model

If we assume all the advantageous mutants in the current population are descended from single copy of the selected mutant, then recombination breaks the chromosomes and mixes the ancestral haplotype with the background haplotypes during the process, creating a “mosaic” pattern along the chromosomes. If we can record the history and trace the origins for every piece of the chromosomes, the SNP positions of haplotypes in the current population can be assigned into two classes: those descended from the ancestral haplotype and IBD (identical by descent) to the ancestral haplotype; or those descended from one of the background haplotypes. For the *i*th haplotype, we can label every SNP position with the two latent states: *S*_*i*,*j*_ ∈ {*AH*, *BH*}, where *AH* stands for “ancestral haplotype” and *BH* stands for “background haplotype”. The states are latent in the HMM as they are unobservable. The transitions between the two adjacent latent states along every chromosome represent the extent of ancestral haplotype sharing, resulting from the joint effects of recombination and hitchhiking. The extent of the ancestral haplotypes is informative for learning the intensity and duration (or allele age) of the selective process.

Consider a single chromosome *i*, and assume for now that we know the mutant position λ for the illustration purpose. In our model, λ is actually a parameter to be estimated by estimating the likelihood ratio scores for each SNP as the putative mutant position along the chromosome. Knowing the mutant position, we divide the SNPs into two groups. We denote the markers to the left “‑1, ‑2,…, ‑ L” and the markers to the right “1, 2,…, R”. where “1” and “‑1” are adjacent to the mutant and so on. The latent states of chromosome i are denoted as the following:

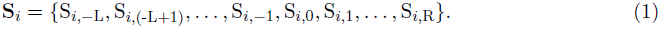

In the following sections, we sometimes simplify the notation by ignoring chromosome subscripts *i* when there is no confusion.

Starting from the mutant position, the two sides of the chromosome form two Markov chains and the time steps of the Markov chains are the SNP positions on the chromosome. The two chains are independent conditioned on the states of the advantageous mutant. Switching between the two latent states is the result of recombination during the selection process. We assume the occurrence of historical recombination along a chromosome follows a Poisson process. The transition matrix of the Markov chains is summarized as follows:

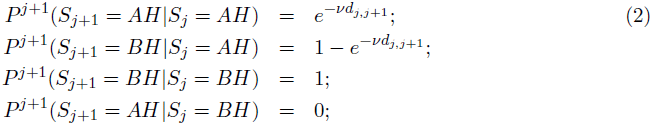

where the *d*_*j*,*j*+1_ is the genetic distance between the *j*th and the (*j* + 1)th SNP, and *ν* is the transition rate per genetic distance, which is directly related to Eqn. (11) in Section 2.5.

The model described above can be used to estimate the probability of being IBD with the ancestral haplotype. But the latent states are hidden and unobserved. Instead, we observe the two alleles of SNPs. If a position is IBD to the ancestral haplotype, and we assume no mutation, the SNP allele, *X*_*i*,*j*_, in that position will be the same as the ancestral haplotype, *A*_*j*_. If mutation is taken into account, the conditional probability is

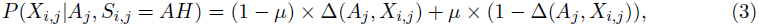

where *μ* is the “mutation rate” for a single SNP, and Δ is the indicator function:

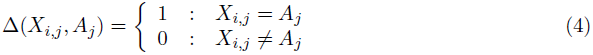

If a locus is IBD to a background haplotype, we can use two distributions to describe the probability of observing allele 1 at that locus. First, a simple binomial distribution is adopted. We use *P*(*X*_*i*,*j*_ = 1|*S*_*i*,*j*_ = *BH*) = *P*_*j*_ (1) to denote the probability of observing allele 1 at the *j*th position, and *P_j_* (1) can be learned from data by the EM algorithm (Durbin et al., 1998). Linkage disequilibrium or the correlation between adjacent loci is not considered in this approach. Second, we model the dependence between adjacent loci using a first order Markov chain (see Zheng and McPeek 2004; Tang et al. 2006 for the applications of the first order Markov chains in modelling linkage disequilibrium). The probability *P*(*X*_*i*,*j*_ = 1|*S*_*i*,*j*_ = *BH,X*_*i*, *j*-1_) needs to be estimated from all haplotypes in the background haplotype groups. So far, we have obtained the probability of observing an allele, *x*_*i*,*j*_, at a locus given its latent state, *s*_*i*,*j*_, being AH or *BH*, *e*_*x*_*i*, *j*__(*s*_*i*,*j*_) (called emission probability in the HMM literature).

With the above emission probabilities, the probability of observing the data at a single position is obtained by summing over all possible latent states of that locus. The likelihood function for the chain to the right side of the mutant is estimated by recursion, the so called forward algorithm. First, we define *f*_*i*,*j*_(*s*_*i*,*j*_) = *P*(*X*_*i*,1_, *X*_*i*,2_,…, *X*_*i*,*j*_, *S*_*i*,*j*_ = *s*_*i*,*j*_), which is the joint probability for the observed sequence up to *j*th step and the latent state at the *j*th step. It is easy to get the recursion equation by the Markov property (Durbin et al., 1998):

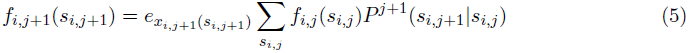

Next, starting from position 0, we set the initial condition as *f*_*i*,0_(*AH*) = *P*(*S*_*i*,0_ = *AH*), which is the probability that the haplotype is descended from the ancestral haplotype. Then by the recursion equation we can get the likelihood function for the right chain of chromosome *i*:

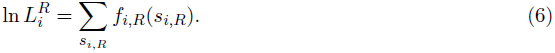

With a similar expression for the left chain, the likelihood function for chromosome i is:

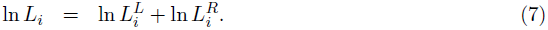

We assume that different chromosomes are independent Markov chains, so we can multiply the likelihood function for each chromosome to form the full likelihood function for the entire data set:

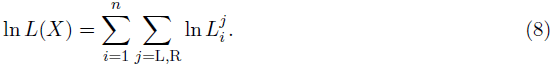

This HMM and the dependencies among variables are graphically illustrated in Fig. 1. Only the right chain of the *i*th chromosome is shown in the figure. The shaded circles denote the observed allele for every SNP position. The hollow circles represent the missing data or unobserved variables, and the directed lines represent probabilistic dependence among these variables.

**Figure 1:**
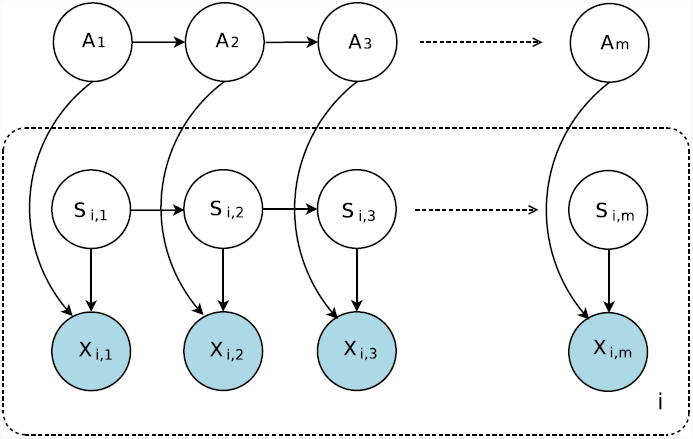
Graphical representation of the hidden Markov model for right chains of the haplotypes from a sample. The first row represents the ancestral haplotype and the dash-lined box is the *i*th haplotype from the sample. The shaded circles denote the observed data, which are the alleles for every SNP position. The hollow circles stand for missing data or unobserved variables. The directed lines represent the probability dependence between the two variables.

### 2.3. Optimization

The ancestral haplotype, **A** = {A_−*L*_,…, A_−1_, A_1_,…, A_*R*_}, is an unknown parameter which needs to be reconstructed when maximizing the likelihood function over parameters. One possible scheme is to exhaustively search over the space of all the possible haplotypes. This is infeasible for large genomic regions since the possible number of haplotypes grows exponentially with the number of SNPs. McPeek et al (1999) used a branch-and-bound algorithm to solve a similar problem in fine-scale disease mapping (McPeek and Strahs, 1999). We use a candidate list method. The candidate list method simply provides a candidate list of ancestral haplotypes chosen from the data and enters into the model each one of them as the ancestral haplotype when estimating the likelihood function. We also tried the branch-and-bound algorithm and another dynamic programming scheme, and found that the candidate list method runs fast and can recover the ancestral haplotype efficiently. Provided a candidate ancestral haplotype, we can iteratively update the transition probabilities and emission probabilities (Eqns. (2),(3)) using the routine Baum-Welch/EM algorithm for HMM (see Durbin et al. 1998 for details).

### 2.4. Hypothesis testing

With the ancestral haplotype identified from the HMM, we propose two tests to detect selection. The selected haplotype group under RPS is expected to be more homogeneous than all the other background haplotypes. The null hypothesis of neutrality corresponds to the case that all the haplotypes are in a similar level of homogeneity. Similar to the former section, we use a first order Markov chain to model the linkage disequilibrium of all background haplotypes. As the neutrality and selection hypotheses are not nested (the two are not overlapping at neutrality), we cannot directly apply the asymptotic theory of likelihood ratio tests here. Instead, we adopt two approaches for hypothesis testing:

- **empirical criterion for likelihood ratio scores.** If we know the population history, we can carry out coalescent simulations to generate a set of neutral samples with allele frequency matching that of the selected allele in the real data, apply the HMM to obtain the likelihood and finally use the simulated likelihood ratios to generate the null distribution for hypothesis testing. With the advent of large-sample genomic sequencing data and the development of efficient methods for inferring population history, it is realistic to have fine-scale population history models that can accurately approximate the true history (Gutenkunst et al., 2009; Gravel et al., 2011; Lukić and Hey, 2012). In genome wide data analysis, we can also use the empirical distribution of the likelihood ratio scores from the genome scan, and pick the top signals (Nielsen et al., 2005; Voight et al., 2006; Pickrell et al., 2009; Chen et al., 2010).
- **permutation test.** Samples are randomly generated by permutating the mutant alleles of haplotypes for M times to allow for the shuffling of haplotype structure linked to the mutant locus. For each of the M samples, the HMM is used to estimate the likelihood ratio, which is used to generate the null distribution for the test. Since computation is intensive for simulating large number of samples by permutation, this approach is feasible only for analyzing data from a local region. And also note that, during the permutation, we only shuffle the mutant alleles or the labels of each haplotype (belonging to the selected or neutral haplotype groups), and thus the linkage disequilibria among marker loci are not broken.

### 2.5. A population genetic model for recent positive selection

The hidden Markov model presented in Section 2.2 is used to identify the ancestral haplotype and detect the breaking points of the ancestral haplotypes retained around each putative selected mutant. The identified ancestral haplotype extent can not only be used to detect the existence of selection, but also to infer the parameters, such as, the section intensity *s* and the duration of the selective process *t*. In this section, we describe the population genetic model needed for parameter inference, and in the next section, we show how the model can be used to infer the two parameters.

Consider a selective sweep that starts with a single copy of the selected allele (a.k.a. a hard sweep, Hermisson and Pennings 2005). Assume that the selected locus has two alleles A and a, with A being the selected allele. Let *y*_*t*_ be the allele frequency of the selected allele A at time *t*. The frequency trajectory of A over time is random. But when selection is strong enough, we can ignore the randomness of the allele frequency trajectories at the very beginning stage of the selection, and thus the trajectory *y*_*t*_ can be approximated by a deterministic curve (Stephan et al., 1992; Braverman et al., 1995). The deterministic process of the selected mutant and the hitchhiking effect were well studied (Maynard Smith and Haigh, 1974; Ohta and Kimura, 1975; Stephan et al., 1992; Durrett and Schweinsberg, 2004). We demonstrate here how the classic theory can be adopted to obtain the allele frequency of the advantageous mutant as a function of the selection intensity and allele age, and the probability distribution of ancestral haplotype extents during the selective process.

Assume an additive model for selection, and let the selective advantage of the allele A be *s*. The change of allele frequency in one generation follows the logistic differential equation (Ohta and Kimura, 1975):

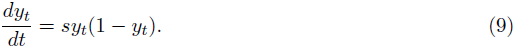

This logistic equation has a solution:

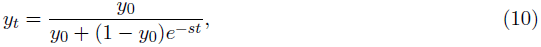

with *y*_0_ being the initial allele frequency of the mutant in generation 0, which can be chosen to be a small value, such as, 1/*N*_*e*_, for a hard sweep process (Kaplan et al., 1988; Stephan et al., 1992).

Now we show how to obtain the distribution of the extent of ancestral haplotypes. Consider a continuous segment between the mutant and a biallelic neutral marker. We use [*AB*] to denote a segment of ancestral haplotype with hidden states A and B being the end points, and [*A*‑] a segment with hidden state A at one end while the other end point of the ancestral fragment arbitrary. Let *P*_*B|A*_ (*t*) be the population frequency of fragment [*AB*] among the A haplotypes at time *t*. Assume that A first appears as on a single haplotype [*AB*] in the population, and all the other haplotypes are |*ab|*, that is, the two loci are in absolute linkage disequilibrium. Note that both A and B take value of *“AH*” in this context. According to Ohta and Kimura (1975), the proportions of chromosomes segments [*AB*] in the “A” group and [aB] in the “a” group at time *t* are respectively (Ohta and Kimura, 1975):

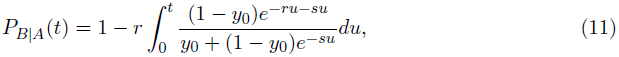

and

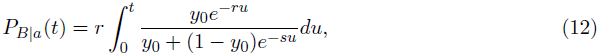

where *r* is the recombination fraction between two loci (see Ohta and Kimura (1975) for details). The above two equations provide the probability distribution for a retained ancestral haplotype with A/a and B/b be the two ends at any time *t* in the selective sweep process. But since the computation involves integration without analytical form we use the following simple approximation. For a random ancestral haplotype [*AB*], if it recombines with any [*A*‑] haplotype during the duration [0, *t*], it does not change the population frequency *P*_*B*|*A*_ (*t*). The only possible change comes from recombination with a haplotype from the neutral haplotype group. The expected number of effective crosses between haplotype [*AB*] and any [*a*‑] haplotype is (Durrett and Schweinsberg, 2004; Chen and Slatkin, 2013):

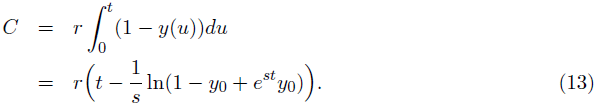

Assume that the number of effective crossovers occurring to ancestral haplotype [*AB*] during the time interval [0, *t*] follows a Poisson distribution. Then the probability of no effective recombination between the [*AB*] haplotype and any [*ab*] haplotypes is

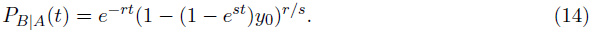

Eqn. (14) provides a very accurate approximation for Eqn. (11) for a wide range of *r* and s values. For example, when *y*_0_ = 0.0001, *t* = 1200, *r* = 0.001 and *s* = 0.01, *P*_*B*|*A*_(*t*) is 0.5926 from Eqn. (11), and is 0.5995 from Eqn. (14).

Let Δr_*R*_ be the recombination fraction between locus *B*_*R*_ and *B*_*R*+ 1_, the probability for the break point occurring at position *B*_*R*_ is

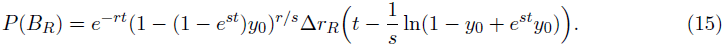

Note that in Eqn. (14) and (15), when st is small, either the selective process is at the early stage or the selection intensity is weak, the term 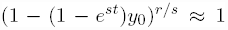, and thus similar to the case under neutrality (Voight et al., 2006; Chen and Slatkin, 2013). However, if *st* is large, the term cannot be ignored. For example, for the same parameter setting as above (*r* = 0.001, *s* = 0.01, *y*_0_ = 0.0001, and *t* = 1200), methods which ignore the term, such as the one presented in Voight et al (2006), can cause a relative bias ≈ 18% (Voight et al., 2006).

### 2.6. Parameter estimation

The ancestral haplotype extents identified by the HMM can be used to estimate the parameters by the importance sampling approach in Chen and Slatkin (2013). Here we show how to estimate the two parameters with a simple but computationally more efficient approach, using the population genetic model presented in the former section.

We take the break points for each chromosome carrying the selected allele, {*B*_*j*,*L*_, *B*_*j*,*R*_, 1 ≤ *j* ≤ *n*_*sel*_}, as the input data, where *B*_*j*,*L*_ and *B*_*j*,*R*_ are the left and right ends of the *j*th ancestral haplotype, and *n*_*sel*_ is the number of chromosomes in the sample carrying the selected mutant. Since we assume that recombination occurs along the chromosomes following a Poisson process, the probability for an ancestral haplotype retained between *B*_*j*,*L*_ and *B*_*j*,*R*_ follows Eqn. (15). We then write the likelihood for the *n*_*sel*_ chromosomes as:

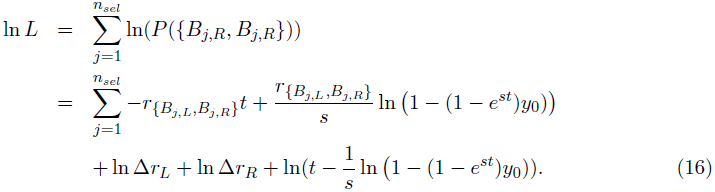

Furthermore, we assume the frequency of the selected mutant in the current generation is known (estimated from the sample, if the sample size is sufficiently large), and a deterministic instead of random frequency trajectory of the mutant (Eqn. (10)), which is reasonable under strong selection. And from Eqn. (10), we obtain the deterministic relation between *s* and *t*:

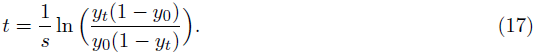

This expression can be substituted into Eqn. (16) to reduce the likelihood as a function of a single parameter *s*. We can easily obtain the estimate 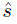, and then get 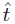 from Eqn. (17).

## 3. Results

### 3.1. Power to detect selection

We used the coalescent simulator *msms* to generate haplotype samples under RPS, and used the samples to evaluate the power of this method in detecting RPS, and the accuracy and precision of selection coefficient estimation (Ewing and Hermisson, 2010). *msms* adopts a structured coalescent scheme to model the effect of a selective sweep on the genealogies of nearby loci. The allele frequency of the mutant at present was chosen to be 0.40 and 0.80, representing selected mutants with moderate and high frequencies. We simulated a sample of 100 haplotypes spanning a region of 1Mb, with the selected mutant located in the middle. The point mutation rate was set to be 1.0 × 10^−8^ per site per generation and the recombination rate 1.0 x 10^−8^. For each level of selection coefficient (2*Ns* = 50,100,200,300, 500,1000), 200 samples were generated. To mimic the effect of phase inference for genotype data, haplotypes were randomly paired to simulate genotype for each individual, and the fastPHASE software was applied to the simulated genotype data to infer haplotypes (Scheet and Stephens, 2006). The output haplotypes were used to evaluate the power of the tests. To give a criterion for hypothesis testing, we generated samples in the neutral case with the same parameters except that the selection coefficient *s* = 0.0. 2000 samples were generated such that the allele frequency of the central SNP was the same as in the selection case. The HMM method was then applied to these neutral samples and the 99th percentile score was recorded as the criterion for significance, which is equivalent to controlling the type-I error to be under 1%. For samples simulated with the given setting (the allele frequency of the selected allele and the selection coefficient), the HMM method was applied, and the likelihood ratio score was recorded and compared to the significance criterion. The proportion of samples with a likelihood ratio score that exceeded the threshold was recorded as the power of the method for the setting.

We compared the performance with two existing popular methods: the allele frequency spectrum-based CLR test by Nielsen et al. (2005), and one long-range haplotype method: the iHS test by Voight et al. (2006). The null distributions for the two statistics were also generated by applying to the simulated data under neutrality, similar to the HMM method.

All the results are plotted in Fig. 2. The haplotype based methods (HMM and iHS) are overall more powerful than the allele frequency spectrum-based method for the two simulated settings (allele frequencies: 0.80 and 0.40), and the difference in performance is more apparent when the selected allele has frequency 0.4. This is not surprising and consistent with the previous conclusion (Sabeti et al., 2002), since the allele frequency spectrum-based method models the hitch-hiking effects of a fixed allele instead of a RPS. The haplotype-based methods, the HMM method and the iHS test, have similar power for both parameter settings.

**Figure 2:**
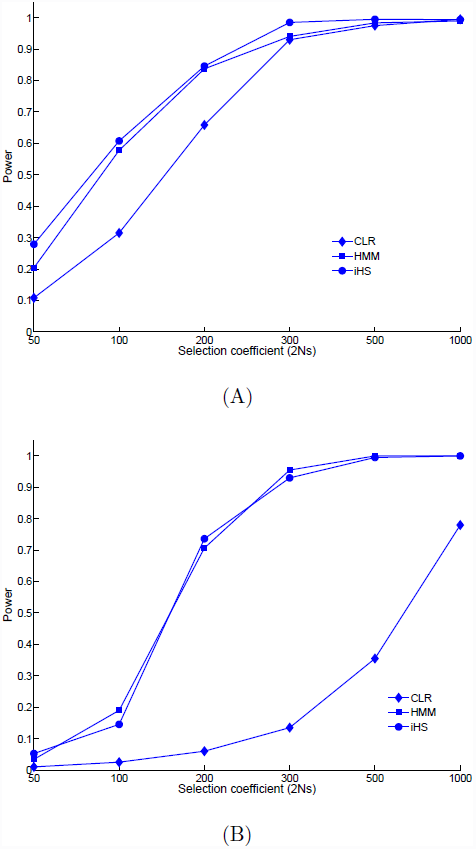
The power of detecting selection for a range of selection intensities. The derived allele frequencies of the selected mutant are (A) 0.80 (B) 0.40. The curve with squares is the proportion of significant results by the likelihood ratio test of the HMM method; the curve with circles is for the iHS test by Voight et al. (2006); and the curve with diamonds is for the composite likelihood ratio test by Nielsen et al. (2005); The 1% cutoff level for the three tests were generated by simulation by assuming the demographic history of the population is known.

### 3.2. Precision and accuracy of the parameter estimation

We also evaluated the accuracy of the parameter estimation. Box plots of the estimated selection coefficients stratified by the true values of selection coefficients from 200 simulations using the above procedures are shown in Fig. 3. The horizontal dashed lines indicate the true selection coefficients in simulation. The bars inside the boxes indicate the medians and the two borders of the box correspond to the first and the third quantiles of the estimates. The medians of the boxes match the true values well, demonstrating that the estimates are accurate. The inter quantile range (IQR) indicates the precision of the estimates for selection coefficients.

**Figure 3:**
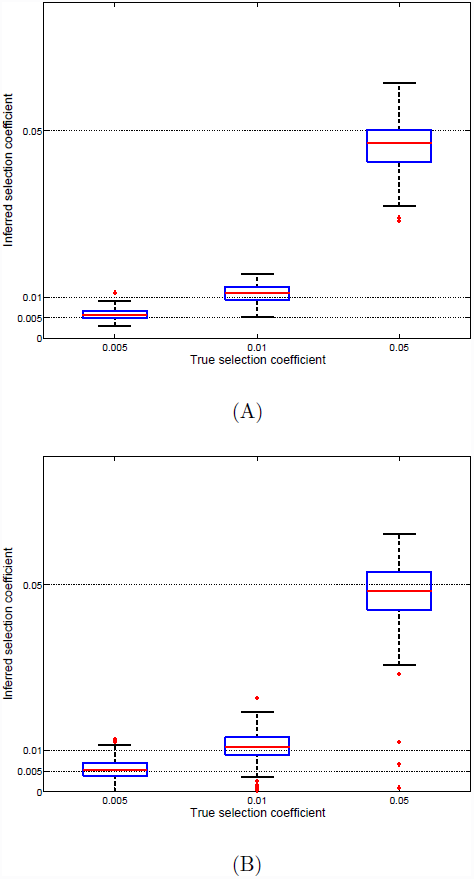
The accuracy of the estimation of selection coefficient for selected mutants with the derived allele frequencies 0.80 (A) and 0.40 (B). The X-axis shows the true values of selection coefficients in the simulations (*s* = 0.005, 0.01, 0.05). The Y-axis shows the inferred selection coefficients. The box plots are estimated values for 200 simulations. Bars inside the box indicate the median of the estimates and the two borders of the boxes correspond to the first and the third quantiles of the estimates.

Box plots of the estimated allele ages stratified by the true values of selection coefficients from 200 simulations are shown in Fig. 4. The inferred allele ages were from applying the HMM method to the simulated data. The true allele ages of the simulated samples were obtained by outputting the allele frequency trajectories during simulations. Since the allele ages vary among simulated samples, we present the log_2_ of the ratio of inferred allele age 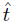 over the true allele age 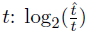. From Fig. 4, we can see the estimates of allele ages for the tested selection coefficient levels *s* = 0.005,0.01 and 0.05 are quite good. In general, the estimates are unbiased, and most of the ratios are within the 2 times and 0.5 times range.

**Figure 4:**
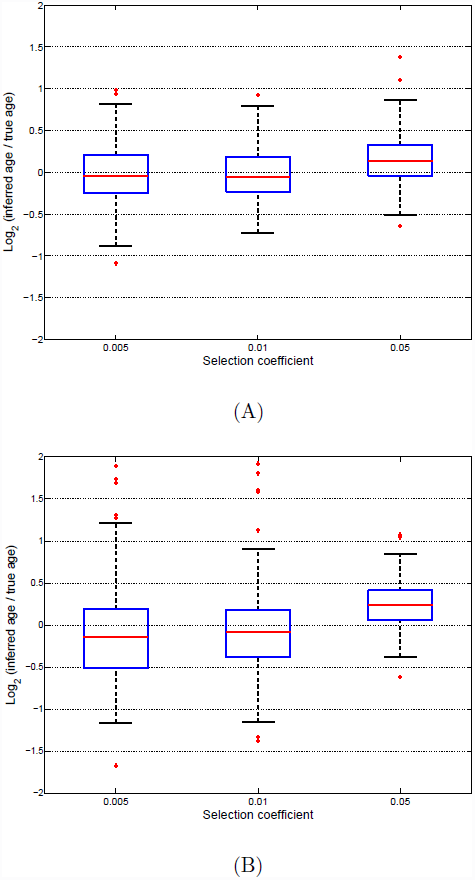
The accuracy of the estimation of allele age for selected mutants with the derived allele frequencies 0.80 (A) and 0.40 (B). The X-axis shows the values of selection coefficients in the simulations (*s* = 0.005, 0.01, 0.05). The Y-axis shows the log_2_ (*inferred age/true age*) values. The box plots are estimated from 200 simulations. Bars inside the box indicate the median of the estimates and the two borders of the boxes correspond to the first and the third quantiles of the estimates.

We compared the performance of the HMM method with two existing methods: the forward simulation method presented in Beleza et al. (2013) (referred to as ForSim in the following paragraphs), and the importance sampling-based method by Chen and Slatkin (2013) (referred to as IS-Age). We applied the two methods to the same simulated data. The inferred selection intensity and allele age were recorded, and the mean and root-mean-square error (RMSE) of the inferred parameters are shown in Tables 1 and 2 for the comparison with the HMM method.

**Table 1:** The comparison of the three methods for estimating selection intensity. The numbers in the table are the mean and RMSE of the inferred selection intensity by the three methods. HMM estimator: the method of this paper; ForSim: the forward simulation method by Beleza et al. (2013); IS-Age: the importance sampling-based method by Chen and Slatkin (2013).

|  | HMM estimator |  | ForSim |  | IS-Age |  |
| --- | --- | --- | --- | --- | --- | --- |
|  | Mean | RMSE | Mean | RMSE | Mean | RMSE |
| freq= 80% |  |  |  |  |  |  |
| $s = 0.005$ | 0.0059 | 0.0015 | 0.0206 | 0.0187 | 0.0055 | 0.0045 |
| $s = 0.01$ | 0.0108 | 0.0022 | 0.0269 | 0.0172 | 0.0114 | 0.0129 |
| $s = 0.05$ | 0.0465 | 0.0070 | 0.0265 | 0.0236 | 0.0381 | 0.0131 |
| freq= 40% |  |  |  |  |  |  |
| $s = 0.005$ | 0.0054 | 0.0026 | 0.0193 | 0.0147 | 0.0075 | 0.0038 |
| $s = 0.01$ | 0.0110 | 0.0039 | 0.0213 | 0.0116 | 0.0127 | 0.0058 |
| $s = 0.05$ | 0.0478 | 0.0086 | 0.0207 | 0.0294 | 0.0325 | 0.0189 |

**Table 2:** The comparison of the three methods for estimating allele age. The numbers in the table are the mean and RMSE of log_2_ (inferred allele age/true allele age) by the three methods. HMM estimator: the method of this paper; ForSim: the forward simulation method by Beleza et al. (2013); IS-Age: the importance sampling-based method by Chen and Slatkin (2013).

|  | HMM estimator |  | ForSim |  | IS-Age |  |
| --- | --- | --- | --- | --- | --- | --- |
|  | Mean | RMSE | Mean | RMSE | Mean | RMSE |
| freq= 80% |  |  |  |  |  |  |
| $s = 0.005$ | -0.0410 | 0.3637 | -1.2113 | 1.2313 | 0.033 | 0.7255 |
| $s = 0.01$ | -0.0190 | 0.3053 | -0.2830 | 0.4125 | -0.0984 | 0.6384 |
| $s = 0.05$ | 0.1469 | 0.3253 | 1.7866 | 1.8039 | 0.2719 | 0.4296 |
| freq= 40% |  |  |  |  |  |  |
| $s = 0.005$ | -0.0450 | 0.7080 | -0.4826 | 0.6284 | -0.3158 | 0.6331 |
| $s = 0.01$ | -0.0621 | 0.5949 | 0.2720 | 0.4275 | -0.1586 | 0.8631 |
| $s = 0.05$ | 0.2884 | 0.6554 | 2.1822 | 2.1994 | 0.4936 | 0.6391 |

Overall, the HMM method outperforms ForSim and IS-Age for the tested parameter range (0.005 ≤ *s* ≤ 0.05, see Tables 1 and 2). The ForSim results show large bias. ForSim only simulates ≤ 8 markers (four on each side of the selected mutant) since the forward simulation is computationally intensive. ForSim adopts a rejection sampling approach to match the real data and simulated data. Instead of using the full sample configuration, the implementation of Beleza et al. (2013) only chooses two summary statistics for the rejection sampling: the allele frequencies of the selected mutants, and the frequencies of the whole intact ancestral haplotypes across all markers. This simplification may explain the bias and big variance of their estimate. However, increasing summary statistics numbers to improve the accuracy may be unrealistic, since the acceptance rate of the rejection sampling becomes extremely low.

IS-Age was expected to performs better than or equally with HMM. The population genetic model for parameter inference in HMM is simplified and approximated with a deterministic process (see Eqn. (9)). When selection is strong (e.g., 0.005 ≤ *s* ≤ 0.05), the approximation is valid, therefore similar results are expected for the two approaches. However, we observed that the results of IS-Age show higher bias and RMSE than HMM for the tested parameter range. It may be due to the following down-sampling procedures when implementing IS-Age. The simulated data was too large to be analyzed by IS-Age, and to implement the analysis, we reduced the size of each simulated sample by randomly choosing 20 haplotypes, and further randomly dropping 50% SNPs. This procedure may reduce sample information and cause potential bias.

In addition to the difference in accuracy and precision of parameter estimation, the computational efficiency of the three methods is also apparent. ForSim of Beleza et al. (2013) uses forward simulation to generate a population of haplotypes, which evolves by generations. The acceptance rate of the rejection algorithm adopted by their method is very low, even when only two summary statistics, instead of the full sample configuration, are used in the rejection sampling. IS-Age uses importance sampling and makes several efficient approximations to reduce the state space of the genealogical process, but it still requires a down-sampling step to reduce sample size and marker numbers. Analyzing a single sample with IS-Age or ForSim typically takes several days. Both ForSim and IS-Age are computationally too intensive, while the HMM method only takes minutes to analyze a sample, and thus is applicable to analyze genome-wide data.

### 3.3. Genome scan on HapMap data

We applied the method to analyze the HapMap Phase III data. The HapMap Phase III data contains genome wide SNP data from 11 world populations. We focused on the three major populations: the Utah residents with Northern and Western European ancestry from the CEPH collection (CEU), the Han Chinese and Japanese populations from East Asia (ASN), and the Yoruban Africans from West Africa (YRI). The phased haplotype data and SNP positions were downloaded from the HapMap Ftp server. The top 20 most significant regions for the three populations from the genome scans are listed in Tables 3-5.

**Table 3:**
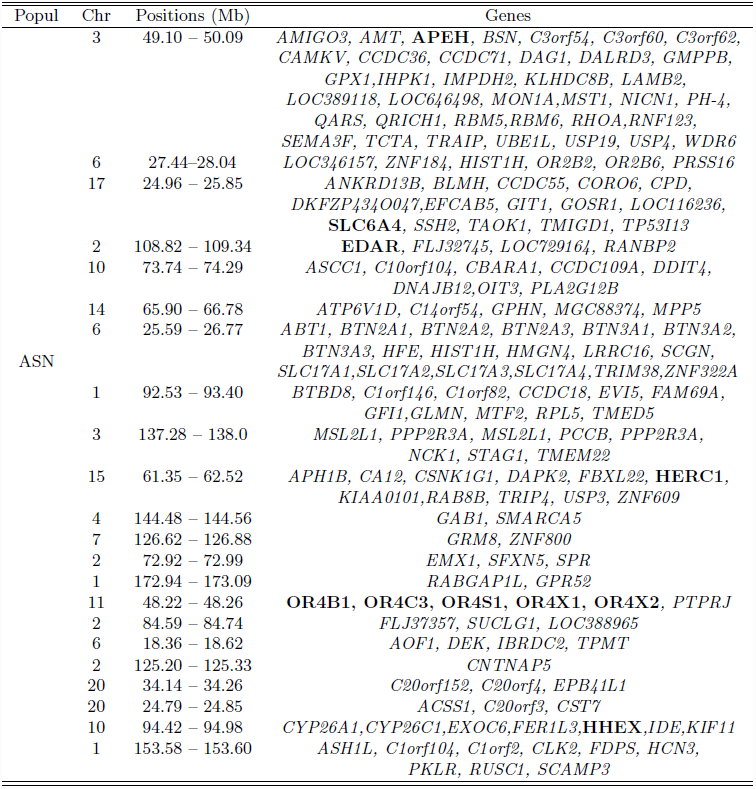
The top 20 regions of the human genome based on the genome-wide scan on the ASN data from HapMap III using the HMM test.

**Table 4:** The top 20 regions of the human genome based on the genome-wide scan on on the CEU data from III using the HMM test.

| Popul | Chr | Positions (Mb) | Genes |
| --- | --- | --- | --- |
| CEU | 6 | 30.0 – 31.68 | *ABCF1, AIF1, APOM, ATP6V1G2, BAT, C6orf134, C6orf136, C6orf15, C6orf205, C6orf47, CCHCR1, CDSN, CSNK2B, DDR1, DHX16, DPCR1, FLJ45422, FLOT1, GNL1, GTF2H4, HCG27, HCG9, HCP5, HLA, IER3, LST1, LTA, LTB, LY6G5B, LY6G5C, MDC1, MICB, MRPS18B, NCR3, NFKBIL1, NRM, POU5F1, PPP1R10, PPP1R11, PRR3, PSORS1C1, RNF39, RPP21, SFTPG, TCF19, TNF, TRIM, TUBB, VARSL, ZNRD1* |
|  | 2 | 135.43 – 136.51 | *ACMSD, CCNT2, CXCR4, DARS, LCT, MCM6, R3HDM1, RAB3GAP1, UBXD2, YSK4, ZRANB3* |
|  | 10 | 73.88 – 74.06 | *CBARA1, CCDC109A, DNAJB12* |
|  | 14 | 66.08 – 66.36 | *GPHN, MGC88374, C14orf54* |
|  | 10 | 74.31 – 74.36 | *CCDC109A, OIT3, P4HA1, PLA2G12B* |
|  | 5 | 130.75 – 131.36 | *FNIP1, RAPGEF6, ACSL6, CSF2, IL3, CDC42SE2* |
|  | 3 | 51.06 – 51.33 | *DOCK3, ARMET, RBM15B, VPRBP* |
|  | 1 | 172.56 – 172.90 | *GPR52, RABGAP1L* |
|  | 1 | 35.94 – 36.03 | *CLSPN, EIF2C4, FLJ38984, PSMB2, EIF2C1* |
|  | 17 | 17.62 – 17.89 | *ATPAF2, C17orf39, DRG2, LRRC48, MYO15A, RAI1, SREBF1, TOM1L2* |
|  | 15 | 41.51 – 41.64 | *ADAL, CATSPER2, CKMT1B, HISPPD2A, LCMT2, MAP1A, STRC, TP53BP1, ZNF690* |
|  | 11 | 48.51 – 49.92 | **OR4A47, OR4C12, OR4C13, FOLH1** |
|  | 4 | 52.66 – 52.69 | *LOC339977, SGCB, SPATA18* |
|  | 15 | 69.95 – 70.28 | *BRUNOL6, GRAMD2, MYO9A, PARP6, PKM2, SENP8, NR2E3, THSD4* |
|  | 2 | 178.06 – 178.26 | *AGPS, PDE11A, TTC30A, TTC30B* |
|  | 5 | 109.97 – 110.01 | **SLC25A46** |
|  | 12 | 87.53 – 87.57 | **KITLG** |
|  | 12 | 110.89 – 111.30 | *C12orf30, ERP29, MAPKAPK5, TMEM116, TRAFD1, PTPN11, RPL6* |
|  | 20 | 33.49 – 34.04 | *C20orf152, PHF20, RBM39, SCAND1, C20orf44, CEP250, GDF5* |
|  | 8 | 51.52 – 51.53 | **SNTG1** |

**Table 5:**
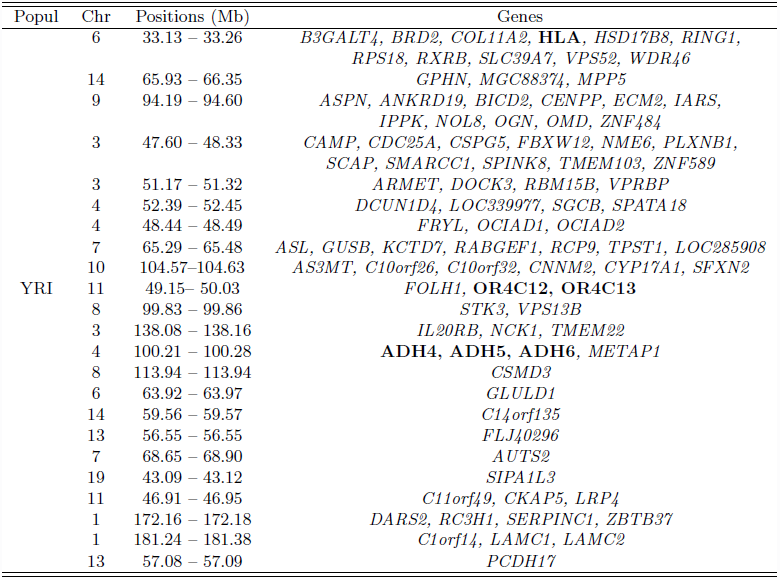
The top 20 regions of the human genome based on the genome-wide scan on on the YRI data from HapMap III using the HMM test.

Some most well-known examples of genes under RPS are among the top lists, and are highlighted in Tables 3-5 with bold type fonts. For example, as shown in Fig. 5, the 135 ‑ 136.5 Mb of Chromosome 2 is one of the most extreme signal in the CEU population. The Lactase gene (*LCT*) is located in the region, and is a famous example of genes under RPS in human populations (Swallow, 2003; Bersaglieri et al., 2004). The ectodysplasin A receptor (*EDAR*) was shown to be one of most significant signals in the ASN population. One non-synonymous mutation of the *EDAR* gene (*EDARV370A*) was absent in other populations, but nearly fixed in the ASN population. Molecular experiments have shown that the mutant is functional for hair thickness and the increase of active eccrine gland numbers, and may contribute to the local adaptation to humid environments in East Asia (Bryk et al., 2008; Kamberov et al., 2013). The other well known examples include pigmentation genes, such as, *KITLG* in CEU; genes related to resistance to infectious disease, such as, *HLA* and *IL3;* skeletal development genes, such as, *GDF5*, and genes related to brain development, such as, *SLC6A4* and *SNTG1.* The extensive identification of well known RPS-target genes demonstrates that the hidden Markov method can efficiently identify the haplotype structure pattern caused by a RPS.

**Figure 5:**
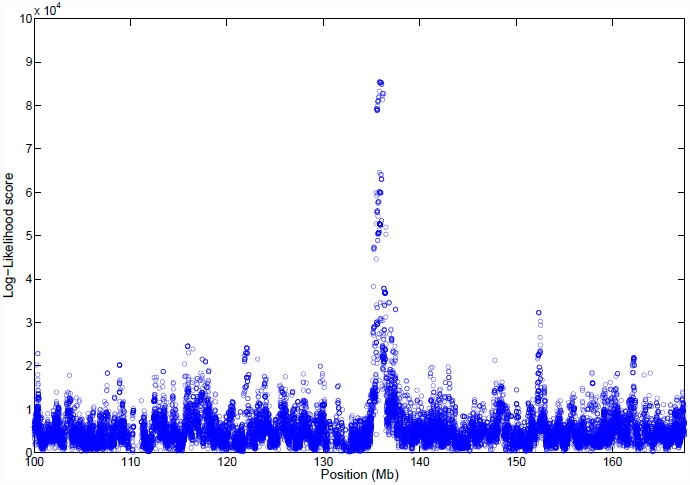
The plot of likelihood ratio scores along chromosome 2 of the Northern European population (CEU) of HapMap III. Lactase gene is located around 136 Mb.

Notice that most of these top regions are identified as targets of RPS by one of former studies (e.g., Akey et al. 2002; Sabeti et al. 2007; McEvoy et al. 2009; Pickrell et al. 2009; Chen et al. 2010; Grossman et al. 2013 etc.). We also carried out a genome-wide analysis using the iHS test by Voight et al. (2006). The iHS scores for all these regions of Tables 3-5 are among top 1% of genome level. This is not surprising, since these most significant regions should be identifiable by various methods. But interestingly, we also found that the ranking of the signals identified by the HMM method is different from that based on the iHS method.

The first example is the 49.10 ‑ 50.09 Mb region of Chromosome 3 in the ASN population. This regions is one of the most significant signals of RPS in ASN genomes, while was not identified as top signals using iHS. We checked the haplotype structure using the HapMap web server and noticed that the ASN haplotypes of this region indeed show very significant haplotype structure related to a RPS (see Fig. S1). There are multiple genes located in this region. Among them, *MST1* (macrophage stimulatory protein 1) is functional for inflammation and wound healing; *APEH (APH)*, a serine peptidase, is known for the “degradation of bacterial peptide breakdown products in the gut to prevent excessive immune response” (Nguyen and Pei, 2005; Raelson et al., 2007). And there are several SNPs identified to be associated with Crohn’s disease by GWAS studies (Raelson et al., 2007). The exact reason for this region being under strong RPS remains unclear, and requests further biological studies. A second example is the gene *SLC25A46.* Its HMM score indicates very significant haplotype structure in CEU caused by a RPS (see Fig. S2). This region was identified as top signals in several other studies based on different approaches, including Chen et al. (2010) and Grossman et al. (2013). The mechanism for it being under RPS is not reported in former literature yet.

### 3.4. Lactase in Northern Europeans

Individuals carrying the lactose tolerance allele in pastoral populations gain some selective advantage for the nutrition provided from dairy. One group of haplotypes in the Northern Europeans has a frequency of ≈ 77%, and extends for > 1Mb. This is consistent with the signals of haplotype structure caused by a RPS. Bersaglieri et al. (2004) used the decay of linkage disequilibrium in the Northern European population to estimate the age of the lactose tolerance associated haplotype to be 2, 538 ‑ 23,954 years ago. They further estimated the selection coefficient of the selected mutant based on the allele age, the allele frequency at present, and a discrete-generation Wright-Fisher model to be 0.014 ‑ 0.15.

We focused our analysis on the 5Mb region of the *LCT* gene from the CEU population of HapMap III (Fig. 5). The genetic distances among SNPs are needed for inferring selection coefficient. We used the Oxford genetic map (Myers et al., 2005), which was also downloaded from the HapMap web page. The genetic position of each SNP was obtained by interpolation. The two SNPs, -13910C/T (rs4988235) and -22018 G/A (rs182549) are believed to be functionally important for lactase persistence in Northern Europeans. We treated rs88235 as the selected mutant when applying the HMM method. We successfully identified the ancestral haplotype in the vicinity of the putative mutant. The posterior probabilities of being at ancestral state for each SNP along the chromosome are inferred. When it is larger than 0.5, the position is labelled as being descended from the ancestral haplotype. The inferred ancestral states of each SNP along the 120 chromosomes are presented in Fig. 6. The ancestral haplotype regions are dark blue and the background haplotypes are cyan. In agreement with former studies, we found the extent of ancestral haplotypes to be more than 1Mb. Some ancestral haplotypes are as large as 2 ‑ 3 Mb. We then used the method in Section 2.6 to estimate the selection coefficient to be 0.0560 (95% CI: 0.0486 ‑ 0.0654), and the age of the selected allele to be 5,350 (95% CI: 4, 580 ‑ 6,163) years, assuming a generation time of 29 years. The 95% confidence intervals were obtained by bootstrapping over haplotypes. This result is consistent with Bersaglieri et al (2004), indicating that *LCT* is one of the genes known to be under the strongest selection effect in humans.

**Figure 6:**
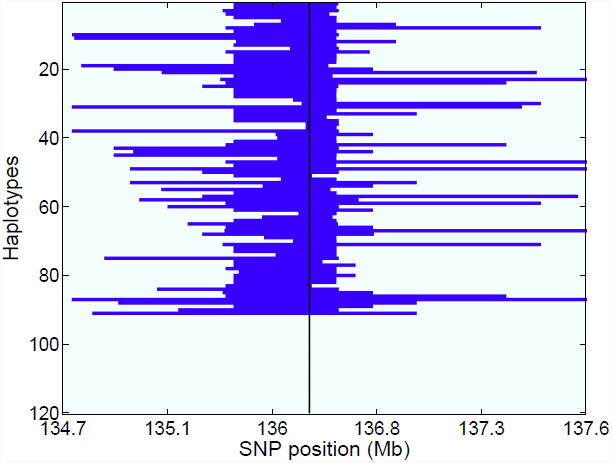
The inferred ancestral states for *LCT* gene region for the Northern European samples from HapMap. Each row represents a single haplotype and each column corresponds to a SNP position. Blue(dark) denotes being inherited by descent from the ancestral haplotype and cyan (light) denotes background haplotypes. The putative mutant is chosen to be ‑13910*C*/*T* (rs4988235), and its position is indicated by the vertical line.

### 3.5. Skin pigmentation genes in Northern Europeans and East Asians

We also investigated three skin pigmentation genes, *KITLG* and *TYRP1* in Northern Europeans, and *OCA2* in East Asians. Light skin color was favoured in Northern Europe and East Asia since it facilitates vitamin D synthesis at higher latitude (Jablonski and Chaplin, 2000). Genes related to the melanosome biogenesis or the melanin bio-synthetic pathways, including *TYR, TYRP1, OCA2/HERC2, SLC45A2, SLC24A5, SLC24A4, MC1R, ASIP, KITLG, IRF4* and *TPCN2*, confer light skin color and show signals of strong RPS (Lao et al., 2007). The evolutionary mechanism of skin pigmentation is at least partially different in Northern Europeans and East Asians. *SLC24A5, KITLG* and *TYRP1* are believed to play important roles in light skin color evolution in Northern Europeans (Lao et al., 2007). In a recent study on pigmentation genes, Beleza et al (2013) used forward simulation and rejection algorithm to get an estimate of the age. They found that selection on *KITLG* happened ≈ 30,000 years ago, after the out-of-Africa migration; and the selective sweep that acted on an European-specific alleles at *TYRP1*, occurred within the last 11, 000 ‑ 19, 000 years, after the first migrations of modern humans into Europe. The mechanism of skin color is less studied in East Asians. A recent association study demonstrated that one non-synonymous polymorphism rs1800414 (His615Arg) of *OCA2* is important for the skin pigmentation in East Asians (Edwards et al., 2010). The time and intensity of selection on the *OCA2* gene in East Asians haven’t been investigated yet.

We applied the HMM method to these three gene regions, which are downloadable from HapMap II. Following the results from Beleza et al(2013), we chose rs642742 for *KITLG* and rs2733831 for *TYRP1* as the putative selected mutants in Northern Europeans. The ancestral states determined by the posterior probabilities of being descended from ancestral haplotypes for each position along the 120 CEU chromosomes are plotted in Fig. 7 and 8. Our estimate of selection coefficient for *KITLG* is 0.0190 (95% CI: 0.0088 ‑ 0.0297), and the allele age is 16, 480 (95% CI:10, 540 ‑ 35, 580) years, assuming a generation time of 29 years. The estimate of selection coefficient for *TYRP1* is 0.0240 (95% CI: 0.0154 ‑ 0.0343), and the allele age is 11, 930 (95% CI:8, 350 ‑ 18, 590) years. Our estimates are roughly consistent with Beleza et al. (2013), and supported the hypothesis that the onset of selection on *TYRP1* may be more recent than *KITLG*, and likely after the split of Eurasian populations. Our estimate of the onset time of selection on *KITLG* is younger than Beleza et al (2013), and does not support the conclusion that it was selected before the separation of Eurasian populations.

**Figure 7:**
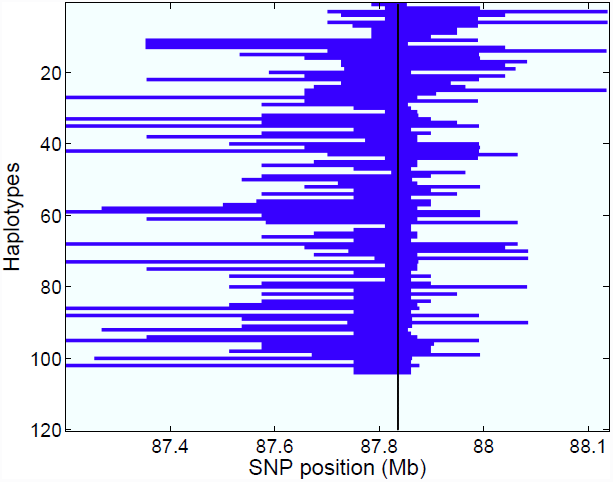
The inferred ancestral states for *KITLG* gene region for the Northern European samples from HapMap. Each row represents a single haplotype and each column corresponds to a SNP position. Blue (dark) denotes being inherited by descent from the ancestral haplotype and cyan (light) denotes background haplotypes. The putative mutant is chosen to be rs642742, and its position is indicated by the vertical line.

**Figure 8:**
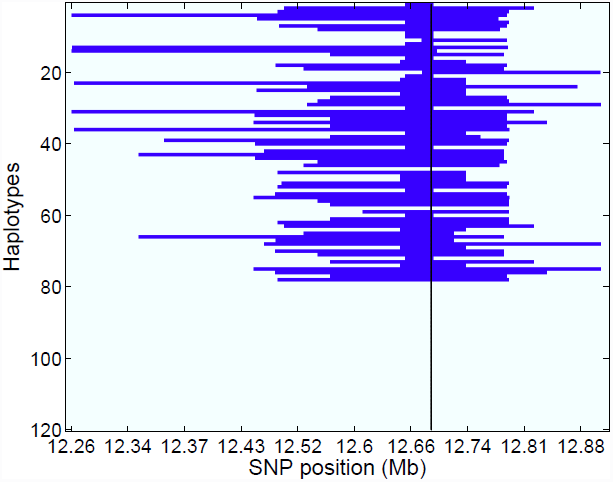
The inferred ancestral states for *TYRP1* gene region for the Northern European samples from HapMap. Each row represents a single haplotype and each column corresponds to a SNP position. Blue(dark) denotes being inherited by descent from the ancestral haplotype and cyan (light) denotes background haplotypes. The putative mutant is chosen to be rs2733831, and its position is indicated by the vertical line..

We also inferred the time and intensity of selection on *OCA2* in East Asians by assuming the non-synonymous polymorphism rs1800414 (His615Arg) as the functional mutant under RPS. The haplotype structure of the *OCA2* region for the 90 Han Chinese haplotypes is presented in Fig. 9. The estimated selection coefficient is 0.0265 (95% CI: 0.0179 ‑ 0.0350), and the allele age is 10,660 (95% CI:8,070 ‑15, 780), which implies that natural selection on *OCA2* in East Asians started independently after the separation of Eurasian populations, and may suggest distinct adaptation mechanisms of skin pigmentation in East Asian and Northern Europeans.

**Figure 9:**
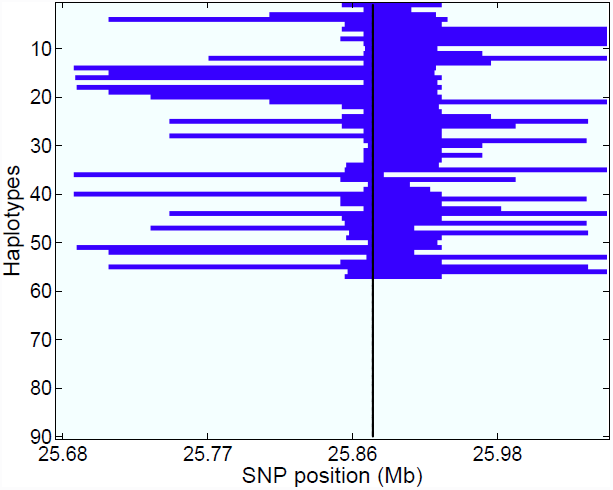
The inferred ancestral states for *OCA2* gene region for the Han Chinese (East Asian) samples from HapMap. Each row represents a single haplotype and each column corresponds to a SNP position. Blue(dark) denotes being inherited by descent from the ancestral haplotype and cyan (light) denotes background haplotypes. The putative mutant is chosen to be rs1800414, and its position is indicated by the vertical line.

## 4. Discussion

We present an HMM method for detecting recent positive selection and inferring selection intensity and allele age when there was a selective sweep. We have shown that the HMM method is effective in capturing the multilocus haplotype structure caused by a RPS. Using coalescent simulations, we showed that the HMM method has more power to detect selection under a range of selection parameters than the allele frequency spectrum-based methods, such as the CLR test (Nielsen et al., 2005). We also demonstrated that its estimates of selection coefficients and allele ages are quite accurate for strong selection. We further illustrated the use of the method by doing a genome scan on the HapMap III data, and analyzing four genes known as the targets of RPS: the *LCT* gene, which conferred to lactose persistence in Northern Europeans, and *KITLG, TYRP1* and *OCA2*, which are related to the skin pigmentation adaptation in Northern Europeans and East Asians. The inferred selection coefficients, ancestral haplotypes and other parameters are consistent with former population genetic studies and human history.

In coalescent likelihood methods, the randomness of trajectories and genealogies are integrated by Monte Carlo methods, which are computationally intensive and only work for investigating a local region (Griffiths and Tavaré, 1994; Kuhner et al., 1995; Chen and Slatkin, 2013). Our method gains computational efficiency by assuming independence among different chromosomes, and constructing the composite likelihood as the product of the marginal likelihood for each chromosome. The performance and the computational efficiency of the HMM method enable its potential applications to genome-wide studies on large-scale data sets. But some simplified assumptions may cause biased estimation when the selection intensity is medium or weak. It is reasonable to believe that if selection is weak or moderate, coalescent-based approaches work better than the HMM method, since ignoring the randomness during early stages of selection can cause bias in estimation (Stephan et al., 1992; Braverman et al., 1995). But on the other hand, the current coalescent-based methods, such as, Chen and Slatkin (2013) are computationally intensive, and are only applicable to a small sample of haplotypes of a local region. A hybrid of the HMM framework and coalescent models, and making use of analytical forms of coalescent distributions to imrpove the computational efficiency (e.g., Griffiths 1984; Chen and Chen 2013), may be a potential direction for further improvement.

The HMM presented in this paper has a potential to incorporate more complex scenarios. For example, it can be used to detect soft sweeps on standing variations, which contain two or more ancestral haplotypes (Hermisson and Pennings, 2005). Another useful extension is to model the haplotype structure in multiple populations.

Note that the proposed method is designed for recent positive selection. If selection is ancient and the mutant has already been fixed in the population, the proposed method will not be useful. For alleles fixed at an earlier time, allele frequency spectrum-based methods, such as Chen (2012), may be a better choice.

## 5. Acknowledgements

We are grateful to Drs. Kun Chen, Thomas Mailund, Noah Rosenberg, and two anonymous reviewers for their insightful comments on an earlier version of the manuscript. We are grateful to Drs. Antόnio Santos and Jorge Rocha for providing their source code and the guidance on using it, to Jared Knoblauch for the assistance in simulation and data analysis. This research was supported by NIH grants R01-GM40282 (to MS) and R01-GM07820 (to JH), and was supported in part by the National Science Foundation through major research instrumentation grant number CNS-09-58854.

---

**Figure S1:**
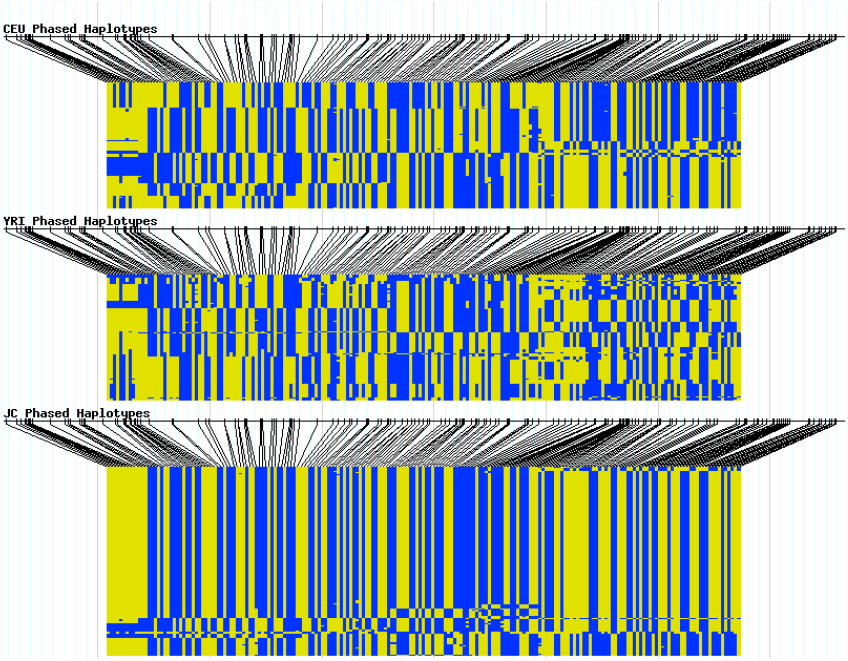
The haplotype structure for the *APEH* gene region for the ASN, CEU and YRI samples from HapMap. Each row denotes a haplotype, and each column corresponds to a SNP. The two colors are for two alleles.

**Figure S2:**
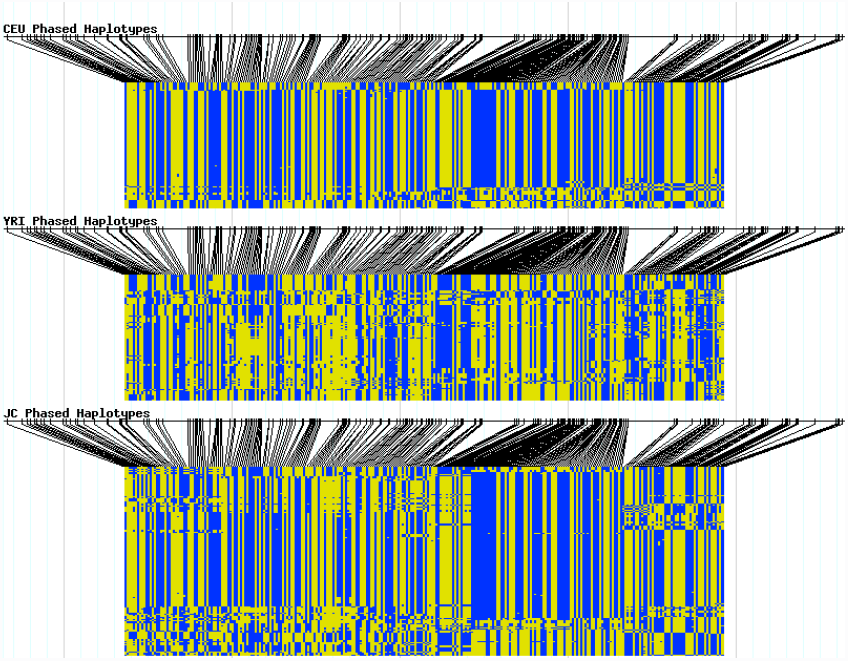
The haplotype structure for the *SLC25A46* gene region for the ASN, CEU and YRI samples from HapMap. Each row denotes a haplotype, and each column corresponds to a SNP. The two colors are for two alleles.

